# Inhibition of the NLRP3 inflammasome prevents ovarian aging

**DOI:** 10.1101/2020.04.26.062646

**Authors:** José M. Navarro-Pando, Elísabet Alcocer-Gómez, Beatriz Castejón-Vega, Jordi Muntané, Pedro Bullon, Chun Wang, Hal M. Hoffman, Alberto Sanz, Gabriel Mbalaviele, Bernhard Ryffel, Mario D. Cordero

**Affiliations:** Cátedra de Reproducción y Genética Humana del Instituto para el Estudio de la Biología de la Reproducción Humana (INEBIR)-Universidad Europea del Atlántico (UNEATLANTICO)-Fundación Universitaria Iberoamericana (FUNIBER); Departamento de Psicología Experimental, Facultad de Psicología, Universidad de Sevilla, Seville Spain; Research Laboratory, Oral Medicine Department, University of Sevilla, Sevilla, Spain; Department of General Surgery, Hospital Universitario Virgen del Rocio/CSIC/Universidad de Sevilla, Sevilla, Spain; Division of Bone and Mineral Diseases, Washington University School of Medicine, St. Louis, MO 63110, USA; Division of Pediatric Allergy, Immunology, and Rheumatology, Rady Children’s Hospital and University of California at San Diego, La Jolla, CA, 92093, USA; Institute of Molecular, Cell and Systems Biology, University of Glasgow, Glasgow G12 8QQ, UK; Laboratory of Experimental and Molecular Immunology and Neurogenetics (INEM), UMR 7355 CNRS-University of Orleans, Orléans, France; IDM, University of Cape Town, South Africa

## Abstract

Inflammation is a hallmark of many processes during aging and reproductive physiology, negatively affecting female fertility. The goal of this study was to evaluate the role of the NLRP3 inflammasome in ovarian aging and female fertility. Age-dependent increased expression of NLRP3 in the ovary was observed in female WT mice during reproductive aging. High expression of NLRP3, caspase 1 and IL-1β was also observed in granulosa cells from patients with primary ovarian insufficiency. Ablation of the NLRP3 inflammasome improved the survival and pregnancy rates in mice, increased hormonal levels of AMH, a biochemical marker of ovarian reserve, and autophagy rates in ovarian tissue. Deficiency of the NLRP3 inflammasome also reduced serum FSH and estradiol levels. Consistent with these results, pharmacological inhibition of NLRP3 using a direct NLRP3 inhibitor, MCC950, improved fertility in female mice to levels comparable to those of *Nlrp3^−/−^* mice. These results suggest that the NLRP3 inflammasome is implicated in the age-dependent loss of female fertility and position this inflammasome as a potential new therapeutic target for the treatment of infertility.

---

Aging is a natural process in all animals involving a progressive impairment of physiological and metabolic homeostasis characterized by many changes in body composition, insulin resistance, mitochondrial and autophagy dysfunction, inflammation and hormonal dysregulation (1). From a clinical point of view, aging is associated with many signs of illness such as cardiovascular, neurodegenerative and metabolic disorders, which are known as age-dependent diseases. These pathologies can be managed by different strategies including pharmacological intervention, lifestyle modifications, and prevention of harmful environmental exposures. However, the ovarian aging process is currently a pharmacologically uncontrollable process that impairs female fertility; the infertility correlates with a rapid decline after age 35, and is attributable to the impairment of the quantity and/or the quality of oocytes (2). Female infertility is exacerbated by socioeconomic changes in developed countries, which enable women to progressively delay the age at which they have their first child. As a consequence, there is increased demand for the treatment of infertility (3).

Inflammation is associated with numerous processes including mitochondrial, oxidative stress and uncontrolled inflammation can negatively affect female fertility (4). However, the underlying molecular mechanisms are poorly understood. Preliminary studies in animal models have shown that genetic deletion of tumour necrosis factor (TNF) receptor improves female fertility, and deletion of IL-1α mice have prolonged lifespan of the ovaries (5,6).

The NLRP3 inflammasome complex is one of the most well-studied inflammatory pathways in human and murine diseases (1). Inflammasomes are multiprotein complexes comprising an intracellular sensor, such as NOD-like receptor (NLR), the adapter protein ASC and pro-caspase-1. Sensing or recognition of the ligand by NLR leads to the maturation of caspase-1 and the processing of its substrates, pro-IL-1β and pro-IL-18 into IL-1β and IL-18, respectively. The NLRP3 inflammasomes is activated by a range of danger and stress signals (7), many of which rise during aging (1). Genetic deletion of Nlrp3 in mice has been shown to improve lifespan and health by attenuating multiple age-related degenerative changes such as cardiac aging, insulin sensitivity with glycemic control, and bone loss (8, 9). However, the role of the NLRP3 inflammasome in ovarian aging and female age-related fertility decline has not been studied. Therefore, we sought to determine whether or not genetic deletion of NLRP3 could have an effect on ovarian aging, and potentially prevent the decline of female fertility.

## EXPERIMENTAL PROCEDURES

### Ethical Statements

For the human study written informed consent and the approval of the Research Ethics Committee of the Virgen Macarena University Hospital, Seville, Spain were obtained, according to the principles of the Declaration of Helsinki.

Animal studies were performed in accordance with European Union guidelines (2010/63/EU) and the corresponding Spanish regulations for the use of laboratory animals in chronic experiments (RD 53/2013 on the care of experimental animals). All experiments were approved by the local institutional animal care committee.

### Patients

The study consisted of 20 women from our fertility clinic diagnosed with Primary Ovarian Insufficiency (POI) and 20 healthy women from our oocyte donation program. Inclusion criteria were healthy women younger than 40 years old, without a previous pregnancy at term, an abnormal ovarian reserve test (antral follicle count <5-7 follicles or FSH >10 mIU/mL and E2 > 60 pg/mL on day 3 of the cycle) and previous characterized poor ovarian response stimulation cycle (≤3 oocytes with a conventional stimulation protocol and E2 < 500 pg/ml the day of Human Chorionic Gonadotropin - hCG-was given). Exclusion criteria were a significant ongoing medical problem. Healthy controls were younger than 35 years old, antral follicle count >10 follicles, serum levels of FSH < 10 IU/mL and serum estradiol < 60 pg/mL on day 3 of the cycle and no previous medical history of infertility.

All patients and controls had taken no drugs or vitamin/nutritional supplements during the 3 weeks before to the collection of blood samples. Clinical data were obtained in the fertility clinic of Instituto de la Biología de la Reproducción Humana (INEBIR) following the committee opinion advice of the Sociedad Española de Fertilidad (http://www.sefertilidad.net), European Society of Human Reproduction and Embryology -ESHRE-(https://www.eshre.eu) and the American Society for Reproductive Medicine (https://www.asrm.org).

### Granulosa cells isolation

Follicular granulosa luteal cells (GCs) were obtained after transvaginal ultrasound-guided oocyte retrieval (TUOR) from 20 patients undergoing *in vitro* fertilization (IVF) treatment at Instituto de la Biología de la Reproducción Humana (INEBIR). Ovarian stimulation was accomplished using a short protocol strategy combining gonadotropin releasing hormone antagonist (an-GnRH) short protocol (Cetrorelix acetate, Cetrotide, Merck Serono Europe Limited, Spain), recombinant FSH (Gonal-f 75 UI, Merck Serono Europe Limited, Spain), urinary human menopausal gonadotropin (hMG) (HMG-lepori 75 UI, Angelini Farmacéutica, S.A, Spain) and recombinant hCG (Ovitrelle 250 micrograms, Merck Serono Europe Limited, Spain) or luprorelin acetate (Procrin 1 mg/0,2 ml, AbbVie Spain S.L.U) to initiate final follicle maturation. Preparing IVF-GCs for culture started when the medical team decided to administer hCG-trigger to optimize the final oocyte maturation. Thirty-six hours before TUOR, light paraffin culture oil, sterile filtered (OVOIL™ - CULTURE OIL, Vitrolife, Sweden), were aliquoted and placed in an incubator for its pre-equilibration at +37.3°C and 6% CO2. At 24 hours before TUOR, culture medium drops (G-IVF™ PLUS, Vitrolife, Sweden) were prepared in the previously equilibrated oil, this prepared medium will contain the oocytes clusters obtained in the TUOR. While preparing the culture medium, two 8 ml aliquots were prepared each one and an aliquot of 1 ml of the same G-mops™ PLUS (Vitrolife, Sweden) in a conical bottomed of 2 ml (medium in which we collected and washed the cumulus oophorus obtained during the TUOR from the remains of follicular fluid and red blood cells) and cultivated in the incubator at 37.5°C and 6% CO2, in that case the tubes must be completely closed since it is an unbuffered medium and would undergo changes in pH and osmolarity if they were left open. The day of the TUOR, the embryologist team worked on a safety work hood with a heated surface (K-SYSTEMS IVF Workstations, CooperSurgical Medical Devices), and the following embryo laboratory material were prepared: 60 mm diameter embryo tested plates, sterile glass Pasteur pipettes, and 10 ml embryo tested conical tubes. During TOUR, the gynecologist aspirated the follicular fluid of each follicle and deposited it in a sterile conical tubes previously heated in a thermoblock which were transferred to the embryo laboratory in a thermoblock to ensure them a constant temperature once the oocytes were aspirated. Simultaneously, the embryologist receives the tubes from the operating room and their contents were deposited in the 60 mm preheated plates in the safety work hood. Later, the cumulus-oocytes were identified under the microscope and GCs were manually collected and deposited in a small conical tube containing 1 mL of MOPS buffered medium containing human serum albumin (G-MOPS PLUS, Vitrolife, Sweden) and subsequently centrifuged at 3200 rpm during 5 minutes. After centrifugation, the supernatant was extracted, obtaining a compact pellet of GCs which were stored in a freezer at −80 °C.

### Animals

For all experiments, only female mice were used. Young and old *Nlrp3^−/−^* and *Asc^−/−^* mice (C57BL/6J background) and *WT/Nlrp3^+/+^/Asc^+/+^* littermate controls, weighing 25-30 g, were maintained on a regular 12 h light/dark cycle. All groups had *ad libitum* access to their prescribed diet and water throughout the whole study. Body weight was monitored weekly. Animal rooms were maintained at 20–22°C with 30–70% relative humidity.

For Nlrp3 gain of function experiments, we bred neonatal onset multisystem inflammatory disease mice (NOMID - *Nlrp3^D301NneoR^* Jax #017971) to Lysozyme M (LysM)-Cre *(Lyz2*^tm1(cre)Ifo^/J Jax#004781) to generate mice mutant and WT littermate controls on a C57BL/6J background) constitutively expressing hyperactivated Nlrp3 in myeloid cells. Ovaries were harvested from 2-week-old mice and were immediately frozen in liquid nitrogen. The weights of WT littermate control mice were in the range of 6-7 g, and NOMID mice were in 3-4 g range consistent with previous reports.

For all experiments with NLRP3 inhibitors, female C57/BL6/J mice from Jackson Laboratory were maintained on a regular 12 h light/dark cycle at 20–22°C. Treatments were started at 8 months of age after randomization into four groups. This study was distributed in two sub-studies: vehicle vs MCC950 and vehicle vs 16673-34-0. These groups correspond to the following treatment: i) standard diet with i.p vehicle (saline) treatment (vehicle group) from Teklad Global 14% Protein Rodent Maintenance Diet, Harlan Laboratories (carbohydrate:protein:fat ratio of 48:14:4 percent of kcal); ii) standard diet with MCC950 treatment (MCC950 group); iii) vehicle group; iv) standard diet with 16673-34-0 treatment (16673-34-0 group). MCC950 was administered by i.p. route at 20mg/kg daily and 16673-34-0 by i.p. route at 100mg/kg daily. All groups had *ad libitum* access to their prescribed diet and water throughout the study. Body weight and food intake were monitored weekly.

## Reagents

Monoclonal antibodies specific for Beclin-1 and p62 were purchased from Sigma-Aldrich (Saint Louis, USA). The anti-GAPDH monoclonal antibody was acquired from Calbiochem-Merck Chemicals Ltd. (Nottingham, UK). Finally, anti-Bcl-2, anti-Bax, anti-ATG12 and anti-MAP-LC3 antibodies were obtained from Santa Cruz Biotechnology. Anti-AMH antibody was obtained from Abbexa (Cambridge, UK). NLRP3 inhibitors MCC950 and 16673-34-0 were obtained from Sigma-Aldrich (Saint Louis, USA) and R&D Systems (Minneapolis, USA) respectively. A cocktail of protease inhibitors (Complete™ Protease Inhibitor Cocktail) was purchased from Boehringer Mannheim (Indianapolis, IN). The Immun Star HRP substrate kit was obtained from Bio-Rad Laboratories Inc. (Hercules, CA).

### Real-time quantitative PCR (qPCR)

The expression of the NLRP3 gene was analyzed by SYBR Green qPCR of mRNA extracted from blood mononuclear cells. Total cellular RNA was purified from the cells by the Trisure method (Bioline, London, UK). RNA concentration was determined spectrophotometrically. Contaminating genomic DNA was eliminated by incubation of 1 μg of total RNA from each sample with gDNA wipeout buffer for 5 min at 42 °C (Quantitect Reverse Transcription Kit, Qiagen, Hilden, Germany). RNA samples were subsequently reverse transcribed to cDNA using the QuantiTect Reverse Transcription Kit (Qiagen, Hilden, Germany). PCR amplifications were conducted with primers targeting NLRP3 (NM_004895.4) and beta-actin, used as an internal control. Thermal cycling conditions used were denaturation at 95 °C for 20 s, 40 cycles of priming at 54 °C for 20 s, and elongation at 72 °C for 20 s. All reactions were performed in duplicate, and reaction mixes without RNA were used as negative controls in each run. Absence of contaminating genomic DNA was confirmed by setting up control reactions with RNA that had not been reverse transcribed. Fold changes in the expression of genes of interest were calculated using the ΔΔCt method.

### Mouse longevity study

Wild type and *Nlrp3^−/−^* female mice (C57BL/6J background), weighing 25-30 g, were maintained on a regular 12 h light/dark cycle. Mice were housed in groups of four to eight same-sex littermates under specific pathogen-free conditions. Individuals were monitored daily and weighed monthly but were otherwise left undisturbed until they died. Survival was assessed using male and female mice, and all animals were dead by the time of this report. Kaplan–Meier survival curves were constructed using known birth and death dates, and differences between groups were evaluated using the logrank test. A separate group of female mice were sacrificed at different ages between 4 and 12 months to study ovaries (histology and western blots).

### Breeding

WT, *Nlrp3^−/−^* and Asc^−/−^ female mice were individually caged with a WT male of proven fertility for a mating period of 1mo. Female mice were weighed at the beginning of the mating period and 14 d after mating, followed by daily weighing. Female mice that gained weight (at least 3 g) were considered pregnant and sacrificed prior to delivery. Pregnancy was defined as having at least one fetus at laparotomy. The number of fetuses per female was recorded.

### Glucose Tolerance Test

Glucose tolerance tests were performed by fasting the mice overnight for 16 h and then injecting glucose (1 g/kg), intraperitoneally. Glucose measurements were performed using a Bayer Contour blood glucose meter and test strips.

### Serum AMH Measurement

Blood samples were drawn via the retroorbital sinus, and sera samples were extracted and kept at −20 °C until use. Measurements were performed using ELISA according to the manufacturer’s instructions (Beckman Coulter). The calculated interassay variation (n = 5) was 8%, and the calculated intraassay variation (n = 37) was 3.2%.

### Leptin and adiponectin

Serum levels of leptin and adiponectin were assayed in duplicate using commercial ELISA kits (R&D Systems, Minneapolis, USA).

### Serum biomarkers

Serum levels of glucose, triglycerides, cholesterol, uric acid, aspartate aminotransferase, alanine aminotransferase and creatine kinase were assayed using commercial kits (Randox Laboratories, Antrim, UK).

### Immunoblotting

Western blotting was performed using standard methods. After protein transfer, the membrane was incubated with various primary antibodies diluted at 1:1000, and then with the corresponding secondary antibodies coupled to horseradish peroxidase at a 1:10000 dilution. Ponceau S staining was selected as a loading control. Specific protein complexes were identified using the Immun Star HRP substrate kit (Biorad Laboratories Inc., Hercules, CA, USA).

### Histological study

After anesthesia of mice, ovaries were excised and immediately placed in 10% neutral-buffered formalin. The ovaries were embedded in paraffin, cut into 5-μm sections, and stained with hematoxylin and eosin. The middle section of each ovary was photographed for an unbiased comparison.

### Statistics

All data are shown as means ± SD. After, evaluation of normality using the Shapiro-Wilk test, statistical differences among the different groups were measured using either an unpaired Student t test or 1-way analysis of variance (ANOVA) when appropriate with Tukeys post-hoc test when appropriate. A Wilcoxon’s ram sum test was used to calculate the statistical significance between telomere-length distributions. A P value of ≤0.05 was considered statistically significant. Statistical analyses were performed using Prism software version 5.0a (GraphPad, San Diego, CA). Asterisks in the figures represent the following: *: P ≤0.05; **: P ≤ 0.01; and ***: P ≤ 0.001.

## Results

### The NLRP3 inflammasome is activated during ovarian aging

To evaluate the role of the NLRP3-inflammasome in ovarian aging, we examined ovaries from female mice of different ages. WT C57BL6/J female mice exhibited a significant increase in body weight over a 12-month period (Figure 1A) and a significant decrease in AMH after the 6^th^ month of life (Figure 1B). Interestingly, NLRP3 protein expression was increased from the 4^th^ month alongside the levels of active caspase 1 (p20) and an age-dependent increase of active IL-1β (p17) (Figure 1C and D). NLRP3 expression was inversely correlated with serum AMH levels (Figure 1E). Since autophagic dysfunction has also been linked to aging, we analysed the expression of the components in this pathway. We observed an increase in the expression of LC3-II as well as other proteins involved in clearance pathways such as p62/SQSTM beginning at 6 months (Figure 1C and D).

**Figure 1.**
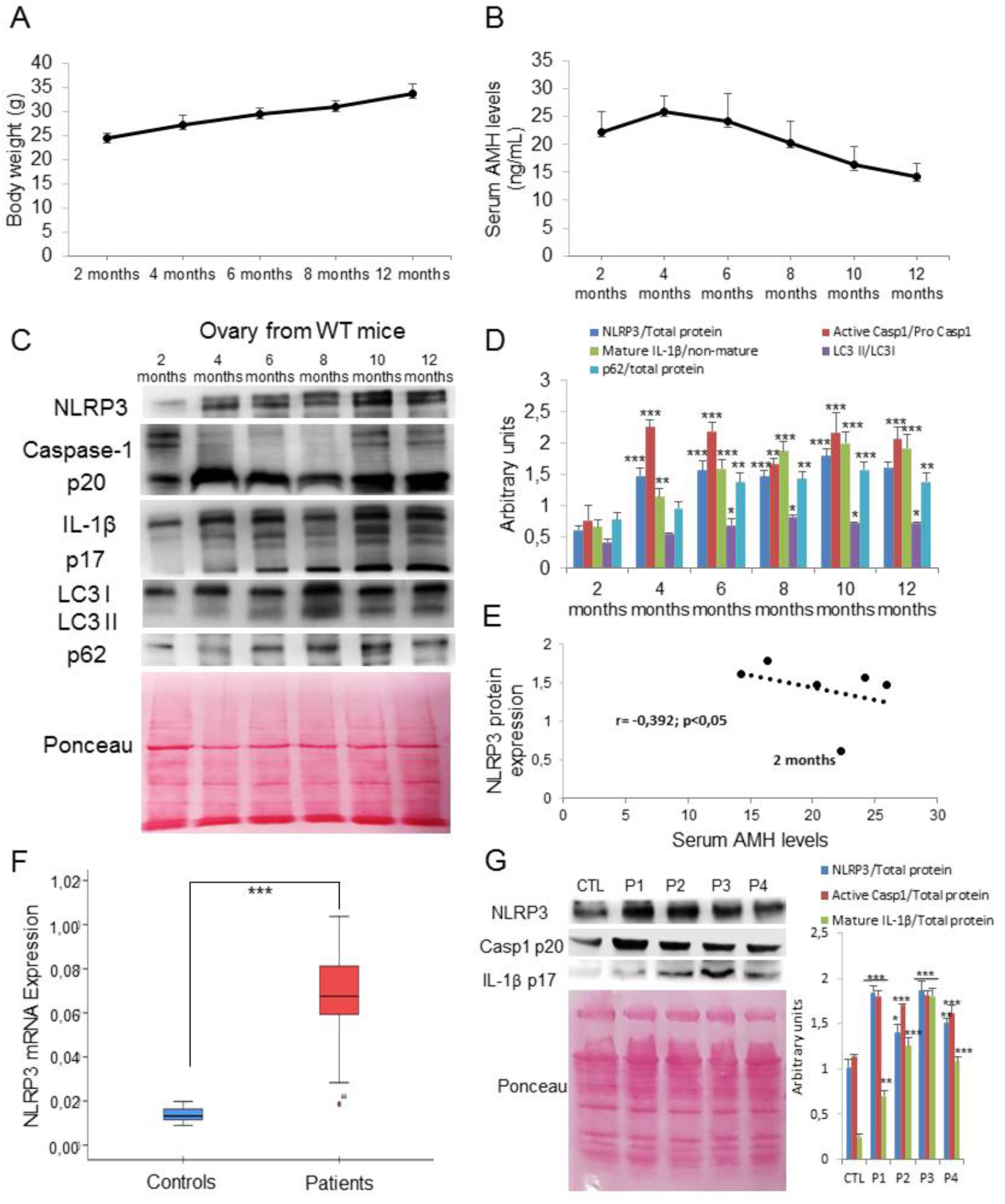
NLRP3 signalling is associated to ovarian aging. (A) Body weights progression of female C57/BL6/J mice evaluated per month. (B) Mean serum AMH levels to evaluate the progression of ovarian reserve during aging. N=8 per group. (C and D) Western blot analysis with representative blots including NLRP3, Caspase 1, IL-1β, LC3, and p62 levels in the ovary of WT mice at different ages. Densitometric analysis is shown as mean ± SD, n = 10 mice; ***P < 0.001, **P < 0.005, *P < 0.05 2mo-old *vs* other ages. N=6-8 per group. (E) Correlation of NLRP3 expression versus serum AMH levels during aging. The correlation was established by calculating correlation coefficients. (F) Human NLRP3 transcript expression levels were determined in granulosa cells by real-time quantitative RT-PCR; n=20 for control and n=20 for POI groups respectively. (G) Western blot analysis with representative blots including NLRP3, Caspase 1 and IL-1β levels in granulosa cells from POI patients compared with controls. Densitometric analysis is shown as means ± SD. ***P < 0.001, **P < 0.005, *P < 0.05 control *vs* patients.

To translate our murine studies to humans, we evaluated NLRP3-inflammasome activation in granulosa cells (GCs) from a human patients with accelerated ovarian aging disease. POI is a critical fertility defect characterized by an anticipated impairment of the follicular reserve for which pathophysiological mechanisms are not understood. Different genetic and molecular alterations have been proposed such as mutations in genes associated with DNA repair or meiosis, bioenergetics alterations and mitochondrial dysfunction (2). The clinical characteristics of POI patients and controls are listed in Table S1 and low ovarian reserve were confirmed by transvaginal ultrasound (Supplementary Figure 1). POI patients showed altered hormonal status compared to the control group (Table S1). FSH levels in these patients were above 10 mUI/mL and E2 levels were above 60 pg/mL. We found significantly increased NLRP3 mRNA and protein expression in GCs from POI patients compared to healthy controls. We also observed elevated protein expression of active caspase 1 (p20) and IL-1β (p17) (Figure 1F and G). However, plasma levels of TNF-α, but not IL-1β and IL-18 were increased in POI patients compared to healthy controls (Table S1).

### NLRP3 expression correlates with reproductive aging

To evaluate the impact of NLRP3 expression on female fertility during aging, we monitored the lifespan and reproductive capacity of *Nlrp3^−/−^* and WT littermates. Kaplan-Meier survival curve showed an increase in mean lifespan of 37% and maximum lifespan of 24% in *Nlrp3^−/−^* (Figure 2A) whereas body weights did not differ between both groups (Figure 2B). Twelve-month-old female *Nlrp3^−/−^* mice exhibited a significant decrease in glucose serum levels at the OGTT peak (>15 min) compared to old WT mice (Supplementary Figure 2A), indicating a higher glucose tolerance consistent with a trend toward lower values of the area under the curve (AUC) of the glucose tolerance test (Supplementary Figure 2B). Further, twelve-month-old female *Nlrp3^−/−^* mice showed no significant difference of serum levels of leptin compared to WT, but an increased levels of adiponectin (Supplementary Figure 2C and D). Twenty eight-month-old WT animals displayed increased age-related alopecia compared to Nlrp3 KO mice (Figure 2C). AMH serum levels were similar in 2-month-old WT and *Nlrp3^−/−^* mice, but were afterward significantly higher in mutant mice compared to WT mice (Figure 2D). Consistent with serum levels, ovarian AMH protein levels were higher in aged (12 month old) *Nlrp3^−/−^* mice compared to WT mice (Figure 2D). Additionally, the levels of the sex hormones E2 and FSH increased between 4 and 12 months in WT mice, but not in *Nlrp3^−/−^* mice (Figure 2E and F). We also analyzed the number of ovarian follicles by histological staining in ovaries from 4 and 12-month *Nlrp3^−/−^* mice and showed more dynamic activity at 12 months and a greater percentage of follicles at 4 and 12 months compared with WT mice. *Nlrp3^−/−^* ovaries retained a larger pool of follicles, which appeared more active in folliculogenesis and contained many corpora lutea, observations suggesting successful ovulation (Figure 2G and H).

**Figure 2.**
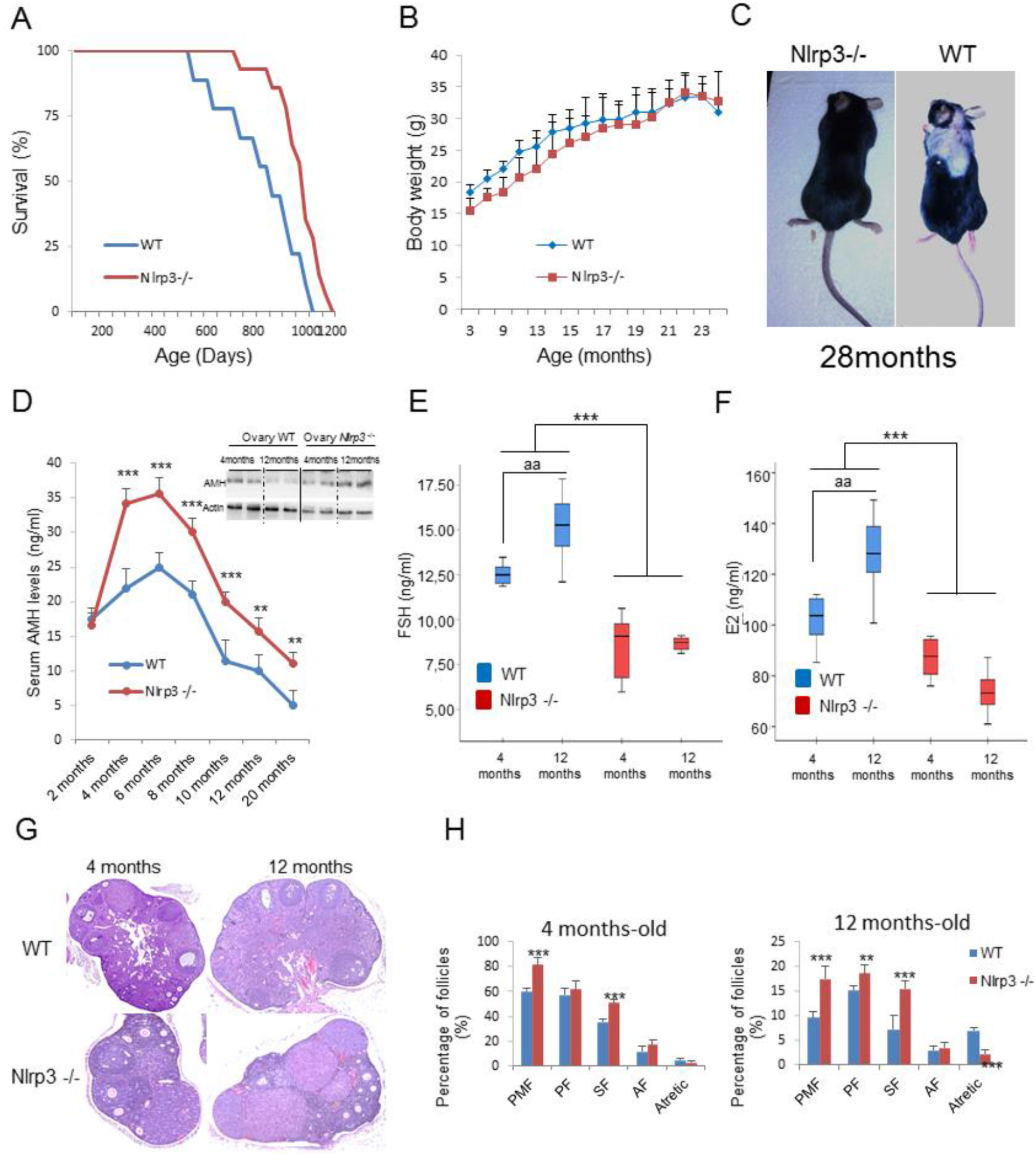
NLRP3 signalling suppression in mice extend lifespan and improve hormonal status in female mice. (A) Kaplan-Meier graph showing a significant increase in n= the maximum lifespan in WT mice compared with Nlrp3−/− mice. (B) Body weights of the groups over time. (C). Representative photographs of 28 months old mice. (D) Mean serum AMH levels by ELISA and ovarian AMH protein levels (top) measured by WB to evaluate the progression of ovarian reserve during aging in WT mice compared with Nlrp3 −/− mice. On top, ovarian AMH protein levels. (E and F) Analysis of serum concentrations of FSH, and E2 measured by ELISA. N=8 per group. (G and H) Representative micrographs of 4 and 12-mo-old WT and Nlrp3−/− ovarian sections. The number of ovarian follicles at developmental stages [PMF, P (primary follicle), SF (secondary follicle), and AF (antral follicle)] was assessed in every fifth serial section of WT and Nlrp3 −/−. N=15-20 per group. Data are shown as means ± SD. ***P < 0.001, Nlrp3 −/− *vs* WT mice; ^aaa^P < 0.001, 4mo-old *vs* 12mo-old mice.

A significant reduction of the pregnancy rate was observed in WT mice during aging but was markedly preserved in aged (12-mo-old) *Nlrp3^−/−^* mice compared with WT mice (Figure 3A). These observations were consistent with AMH and ovary data. Importantly, mean litter size was also significantly higher in 12-month-old *Nlrp3^−/−^* mice compared with littermate mice as in young mice (4 months) (Figure 3B). Interestingly, the loss of the apoptosis-associated speck-like protein containing a CARD (ASC), which has a pivotal role in the assembly of several inflammasomes (7), did not affect the age-dependent loss of female fertility. Indeed, *Asc^−/−^* mice showed similar levels of AMH, FSH and E2 during aging and reduced AMH protein expression (Supplementary Figure 3A-C). The pregnancy rate and mean litter size in *Asc^−/−^* mice were not statistically different compared with WT mice (Supplementary Figure 3D and E). These observations suggest that NLRP3 attenuates the process of ovarian aging independent of ASC expression.

**Figure 3.**
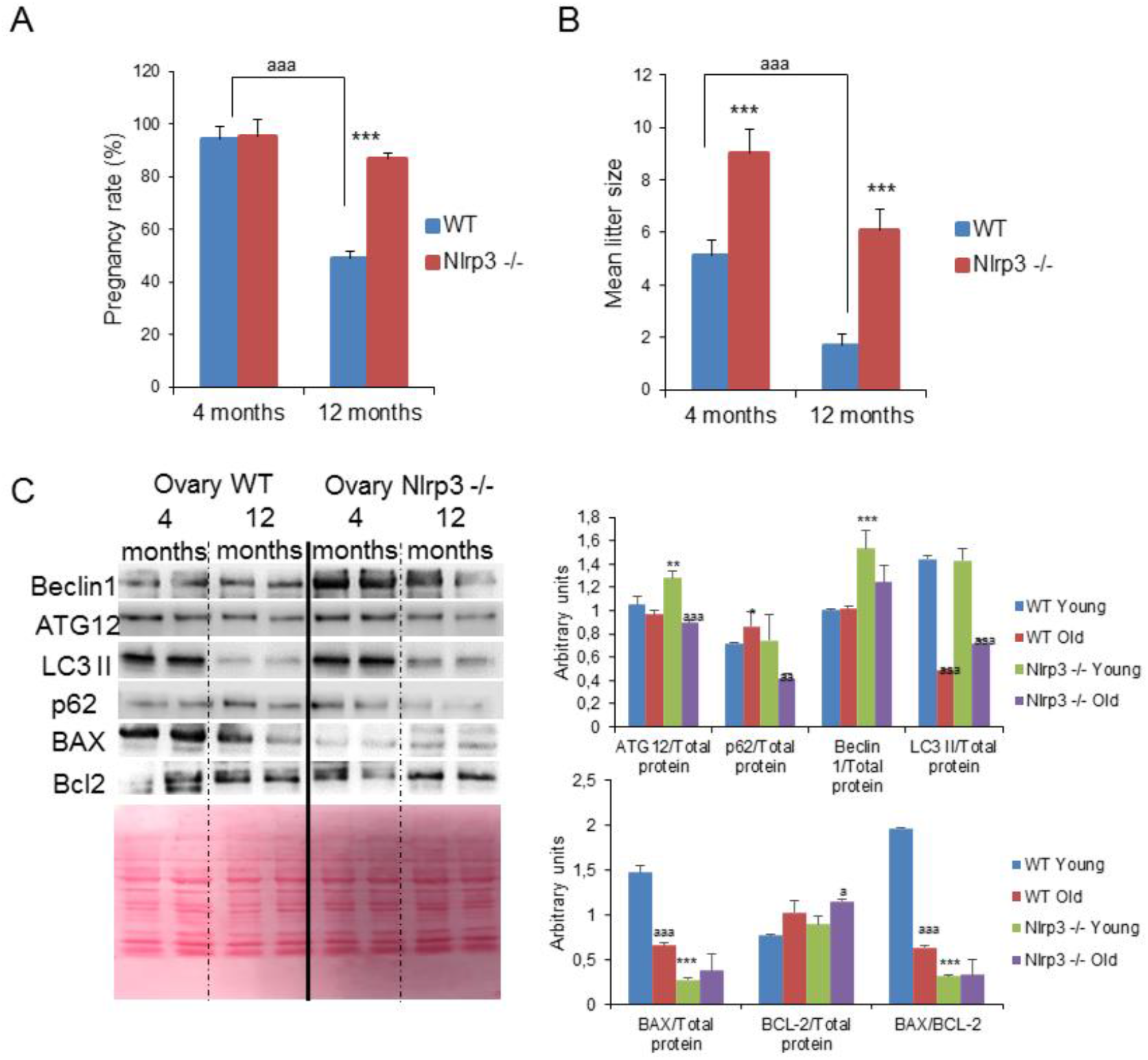
NLRP3 deletion improves reproductive rate and autophagy pathway. Pregnancy rate (A) and mean litter size (B) in aged WT and Nlrp3 −/− mice. Dara are shown as means ± SD. ***P < 0.001, Nlrp3 −/− *vs* WT mice; ^aaa^P < 0.001, 4mo-old *vs* 12mo-old mice. (C) Western blot analysis with representative blot including BECLIN 1, ATG12, LC3, p62, BAX and Bcl2 levels in the ovary of 4 and 12-mo-old WT and Nlrp3 −/−. Densitometric analysis is shown as means ± SD, n = 10 mice per group and age. ***P < 0.001, **P < 0.005, *P < 0.05 WT *vs* Nlrp3 −/−; ^aaa^P < 0.001, young *vs* old mice.

To reinforce the role of NLRP3 in female fertility, we studied the effects of a gain of function NLRP3 mutant associated with neonatal-onset multisystem inflammatory disease (NOMID) in mice (10). We generated Nlrp3fl(D301N)/+; LysM-Cre mice for conditional NLRP3 activation in only myeloid cells. A significant difference of bodyweight was observed between female WT and NOMID mice (6-7 g for 2-week-old WT mice vs.3-4 for NOMID mice; and Supplementary Figure 4A). NOMID female mice showed significantly reduced serum levels of AMH with also reduced ovarian AMH protein levels (Supplementary Figure 4B). E2 and FSH serum levels were also higher in NOMID mice compared to WT littermates; although in the case of E2 did not reach statistical significance (Supplementary Figure 4C and D). These data would likely be consistent with infertility, however these mice die at approximately 3-weeks prior to reproductive age.

### Age-related autophagic and apoptotic changes in the ovaries are prevented by NLRP3 ablation

Blockade of autophagic flux and accumulation of non-degraded substrates in the form of autophagosome are linked to aging (11). Furthermore, apoptosis and autophagy are involved in the process of oocyte loss (12, 13). Interestingly, *Nlrp3^−/−^* female mice showed increased levels of ATG12, beclin 1 and LC3II protein, and reduced expression of p62/SQSTM1 (Figure 3C). Apoptosis has been associated to the ovary aging, also associated to inflammation (6). The expression of the pro-apoptotic proteins Bax was also substantially lower in 12-month-old *Nlrp3^−/−^* ovaries compared with WT ovaries despite no significant changes in the anti-apoptotic protein BCL-2 (Figure 3C). Interestingly, we observed increased levels of TNF, and IL-6 in age 12-month-old *Nlrp3^−/−^* ovaries compared to WT ovaries (Supplementary Figure 5). These results suggest that loss of NLRP3 has a role in the reproductive aging independent of other inflammatory pathways.

### Pharmacological inhibition of NLRP3 rescues female fertility during ageing

We tested the effect of the NLRP3 inhibitor, MCC950 in female mice. Because fertility in mice declines around 8 months of age, we treated 8-month-old mice with vehicle and MCC950 (20mg/kg/day, i.p.) for 12 weeks. No significant changes in body weight were observed between both groups (Figure 4A). Vehicle treated mice showed significantly reduced AMH serum levels compared to MCC950-exposed mice (Figure 4B). Serum levels of FSH and E2 were also decreased by MCC950 treatment (Figure 4C and D), findings that were consistent with histological staining showing higher percentages of follicles and lower percentages of atretic follicles in MCC950-treated compared to vehicle administrated mice (Figure 4E and F). These data revealed a gradual decline in female fertility in vehicle-treated mice whereas age-matched mice in the MCC950-treated group remained fertile. In fact, the pregnancy rate was markedly increased in MCC950-treated mice compared to the vehicle group (Figure 4G), though differences in the mean litter size were not observed (Figure 4H).

**Figure 4.**
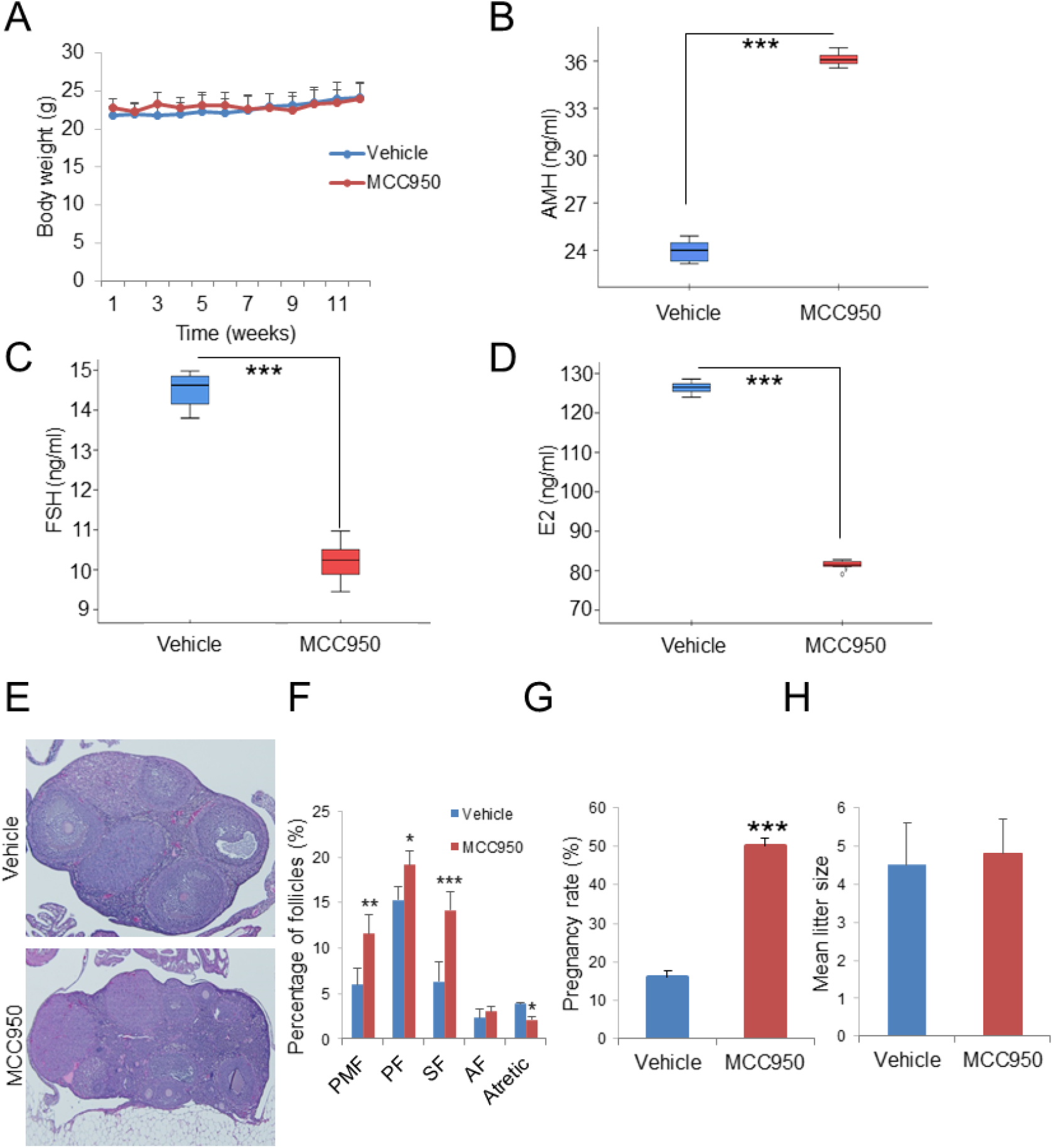
NLRP3 pharmacological inhibition improves female fertility. (A) Body weight chart of 8-month old female mice treated either with vehicle or MCC950. Mice were fed with the standard diet for 12 weeks and their body weights were monitored weekly. Hormonal status in female mice given MCC950 treatment was determined by (B) AMH, (C) FSH and (D) E2 serum levels in female compared with vehicle treated mice. (E and F). N=8 per group. Representative micrographs of vehicle and MCC950 ovarian sections and quantification of the number of ovarian follicles. N=10 per group. Pregnancy rate (G) and mean litter size (H) in vehicle and MCC950 treated mice. N=15 per group. Dara are presented as means ± SD. ***P < 0.001, MCC950 *vs* vehicle treated mice.

In line with data from *Asc^−/−^* mice in which we observed an independent effect of the NLRP3 inhibition, we also analyzed the effects of 16673-34-0, an intermediate substrate produced during glyburide synthesis, which was shown to inhibit NLRP3-inflammasome complex formation by interfering with events involved in NLRP3 conformational changes secondary to activation or binding to ASC (14). Again, 8-month-old female mice were treated with vehicle or 16673-34-0 (100mg/kg/day, i.p.) for 12 weeks. No significant changes were observed in body weight between groups either (Supplementary Figure 6A). Notably, AMH, FSH and E2 serum levels were not significantly different between both groups (Supplementary Figure 6B-D). Finally, there were also no differences in pregnancy rate and mean litter size (Supplementary Figure 6E and F).

## Discussion

Aging of the world population is as undeniable as its profound health and economic consequences (15). The increased average age at which women have children and the consequent increase in problems related with fertility are the principal causes that lead to an increase in the demand for assisted reproduction technologies (16, 17). However, the molecular mechanisms of ovarian aging, which may provide new clues for protection from aging-specific female reproductive decline are still not well understood. Inflammation and autophagy are highly associated with aging and many rejuvenation strategies rely on anti-inflammatory and autophagy inductions (18, 19). Increased systemic inflammation is commonly associated with metabolic alterations, including the appearance of increased adiposity, insulin resistance, and dyslipidemia, responses that are likely critical determinants in the shortening of lifespan and overall health. Many of these metabolic alterations have been associated to NLRP3-inflammassome activation (1, 8–10). However, little is known about the role of NLRP3 during female aging and fertility. Effective fertility requires a fine balance between pro and anti-inflammatory mediators and reproductive hormones. Inflammation has been associated with female infertility and metabolic dysfunction, suggesting that early and chronic inflammation may trigger early menopause, which is associated with the cessation of the ovarian function (20). In the current study, we provide evidence showing that the NLRP3 inflammasome is an important determinant of fertility in human and aged female mice. We show that the expression of NLRP3 and other components of the complex rapidly increase in the ovary during aging, and human GCs, a condition of accelerated ovarian aging. Inflamm-aging, the long-term result of the chronic stimulation of the innate immune system with continuous damage during aging process, has been associated with POI patients in which levels of several cytokines are increased (21). Our data reveal increased levels of TNF-α in serum samples with no significant changes in IL-1β and IL-18. Interestingly, mRNA expression levels of NLRP3 were increased in patients’ GCs. According to this, NLRP3 has a specific role in the ovarian aging process. Indeed, our data also demonstrate that the genetic inhibition of NLRP3, but not in ASC, enhances ovulation and pregnancy rates and improve ovarian endocrine function in an apparently inflammasome-independent manner, whereas ovaries from *Nlrp3* gain of function mutant mice have hormone levels consistent with reduction of fertility. These data suggest that NLRP3 has differential roles in innate immunity and reproduction. Our data are consistent with previous reports showing the preventive role of NLRP3 deletion, but not caspase-1/11 or ASC in other tissues and pathologies (22, 23). Furthermore, *Nlrp3*^−/−^ female mice showed increased inflammatory cytokine levels in the ovary, however, despite this, ovarian aging was prevented by Nlrp3 deletion, which indicates an inflamm-aging-independent role of NLRP3.

Nlrp3 ablation also showed an increase in autophagy in aged ovaries consistent with previous data in other tissues (8, 24). It is known that autophagy efficiency is reduced during aging in many tissues (25). In fact, the autophagy pathway has been linked to the preservation of the oocyte longevity by maintaining the endowment of female germ cells prior to the establishment of primordial follicle pools in the ovary (26, 27). This mechanism may be key to the improvement of ovarian lifespan induced by the inhibition of NLRP3. Further, the support of many of the strategies to improve the reproductive aging and ovarian lifespan through the improvement of autophagy (26, 28, 29).

The present study further demonstrate a metabolic regulatory effect of the NLRP3 inflammasome in females in the reproductive hormone during the equivalent perimenopause period in female mice. Mice are sexually mature by 3–6 months of age and the human endocrine equivalent of human perimenopause is 8-9 months of age (30). During this biological timeframe, many biochemical changes mark reproductive aging, including increased levels of estrogen, FSH, and metabolic and inflammatory factors (31, 32). Our middle-aged *Nlrp3^−/−^* female mice showed the hormonal status of the perimenopause stage, decreased risk factors of age-related diseases. Indeed, female KO mice showed improved glucose tolerance, a non-significant reduction of leptin levels and an increase of adiponectin levels.

Several NLRP3 inhibitors have been identified and studied in different disease states. While several of these inhibitors have been shown to have a specific direct action on NLRP3, others have indirect inhibitory effects (14). MCC950 is a specific small-molecule inhibitor of this inflammasome, although the molecular mechanisms of action of this compound has not been fully elucidated (14). Our study shows that MCC950 improve fertility in female middle-age mice similar to Nlrp3 KO mice.

We acknowledge the limitations of using POI patients only. The use of GCs from healthy woman with different age during the biological aging could corroborate our hypothesis. However, it difficult to obtain samples from healthy individuals at different age for ethical reasons. On the other hand, we have studied the effect of the ablation or constitutive activation of NLRP3 where we cannot separate local out of the systemic effects. Thus, future investigations should focus on ovarian specific ovarian-restricted manipulation of the inflammasomes.

In conclusion, NLRP3 inhibition attenuates ovarian aging and prolongs fertility in female mice. NLRP3 ablation improved hormonal and metabolic characteristics related to female reproductive aging, improved autophagy and reduced apoptosis Therefore, prevention of the ovarian aging process through multiple mechanisms by NLRP3 inhibition could attenuate the reproductive aging in female mice. Taking into account the observed increased expression of NLRP3 in the ovary during normal aging and in POI patients, NLRP3 inhibition offers a promising therapeutic target to maintain female fertility during aging.

## Acknowledgments

This study was supported by a grant from the Andalusian regional government (Grupo de Investigacion Junta de Andalucia CTS113, Consejería de Salud de la Junta de Andalucia: PI-0036-2014) and European funding in Region Centre-Val de Loire (FEDER N° 2016-00110366 BIO-TARGET and EX005756 BIO-TARGET II). GM is supported by NIH/NIAMS AR068972 and AR076758 grants.

