## Supplementary data for "Inhibition of the NLRP3 inflammasome prevents ovarian aging"

**Running Title:** NLRP3 inflammasome and Ovarian Aging

**Corresponding Author**:

Cátedra de Reproducción y Genética Humana del Instituto para el Estudio de la Biología de la Reproducción Humana (INEBIR)-Universidad Europea del Atlántico (UNEATLANTICO)-Fundación Universitaria Iberoamericana (FUNIBER),.

**Tables**

**Table 1. Characteristic of control and POI patient groups**

| | ***Control group*** | ***POI groups*** | ***P value*** |
|---|---|---|---|
| ***Age*** | 31.8±4.1 | 33.1±3.6 | 0.293 |
| ***Sex*** | Female | Female | --- |
| ***Alcohol consumption*** | Sporadic (39%) | Sporadic (42%) | --- |
| ***Tobacco consumption*** | Sporadic (26%) | Sporadic (23%) | --- |
| ***BMI*** | 24.1±4.4 | 24.5±3.2 | 0.744 |
| ***Overweight, BMI 25.0–29.9 kg/m$^2$*** | 9% | 21% | --- |
| ***Obesity BMI 30 kg/m$^2$*** | 9% | 21% | --- |
| ***Estradiol (*** *pg/mL)* | 30.1±2.1 | 71.3±8.1 | 0.0001 |
| ***FSH*** | 4.2±1.3 | 19.5±2.7 | 0.0001 |
| ***TSH*** | 2.1±0.3 | 14.5±1.3 | 0.0001 |
| ***IL-1β*** | 3.7±1.4 | 3.1±1.4 | 0.183 |
| ***IL-18*** | 3.1±2 | 2.8±1.3 | 0.577 |
| ***TNF-α*** | 14.9±5.8 | 50.5±8.7 | 0.0001 |

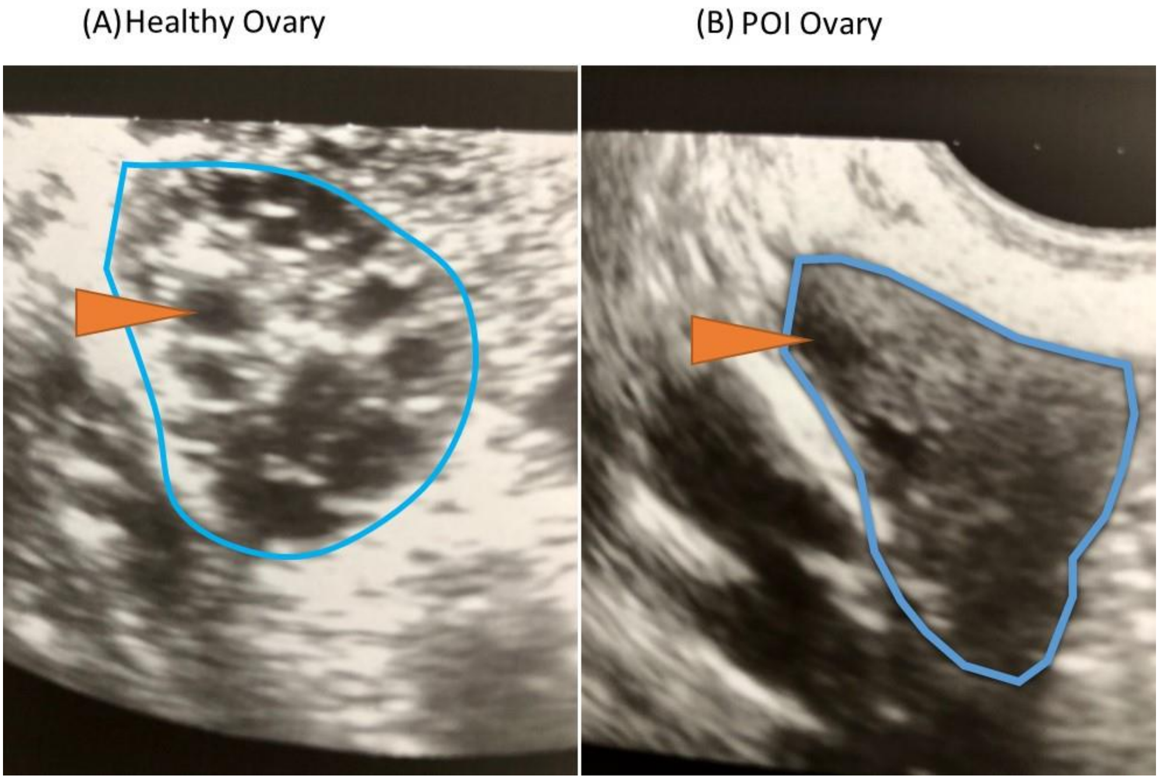


Supplementary Figure 1. The image shows a vaginal ultrasound on the third day of the menstrual cycle with an example of a healthy ovary (A), and one (B) of a characteristic ovary of premature ovarian failure (POI). An arrow indicates the ultrasound appearance of a follicle. It can be clearly seen that the number of follicles in ovary A is greater than in ovary B.

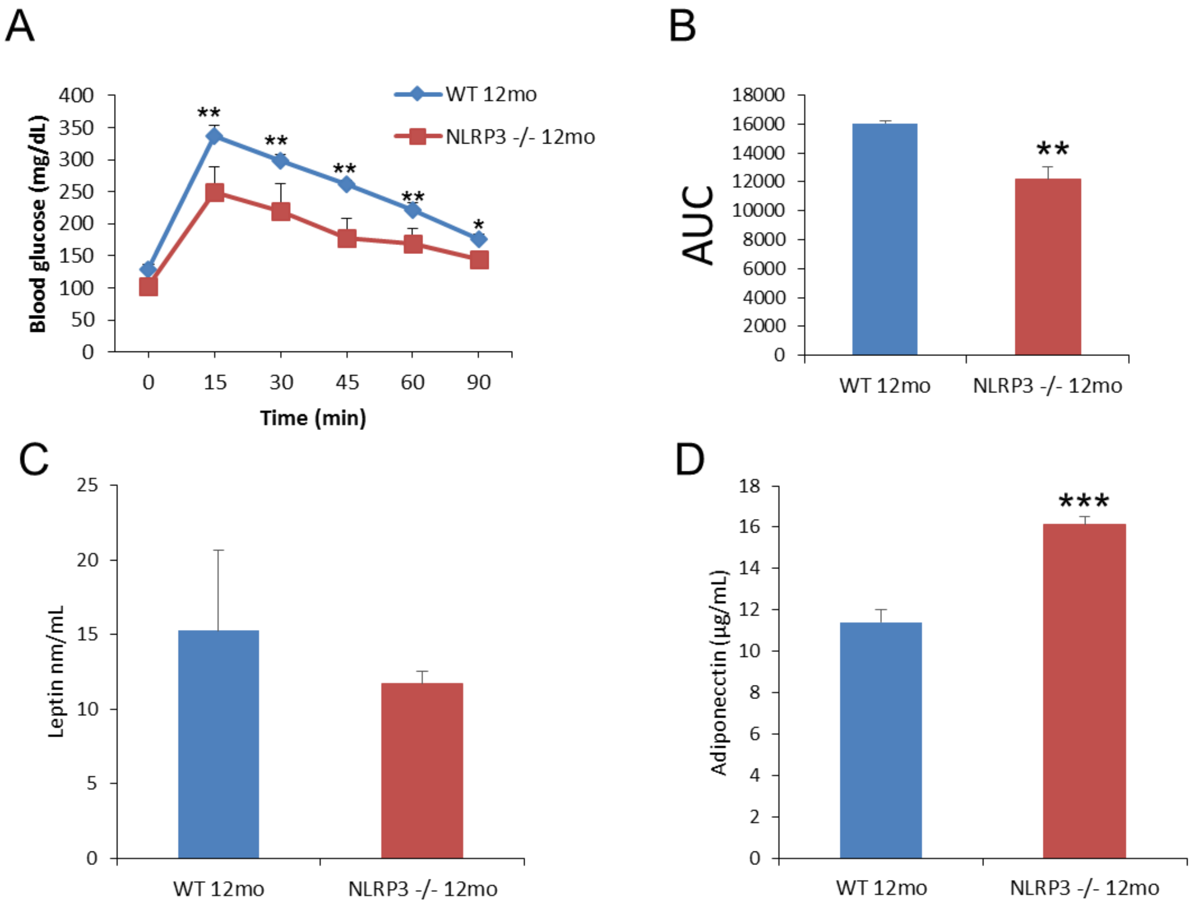


Supplementary Figure 2. (A and B) Oral glucose tolerance test with area under the curve. (A and D) Levels of leptin and adiponectin. Blood samples were collected after overnight fasting. All data are presented as means ± SD, n = 10 mice; *P < 0.05, **P < 0.005, ***P < 0.001 WT *vs* NLRP3 -/- mice.

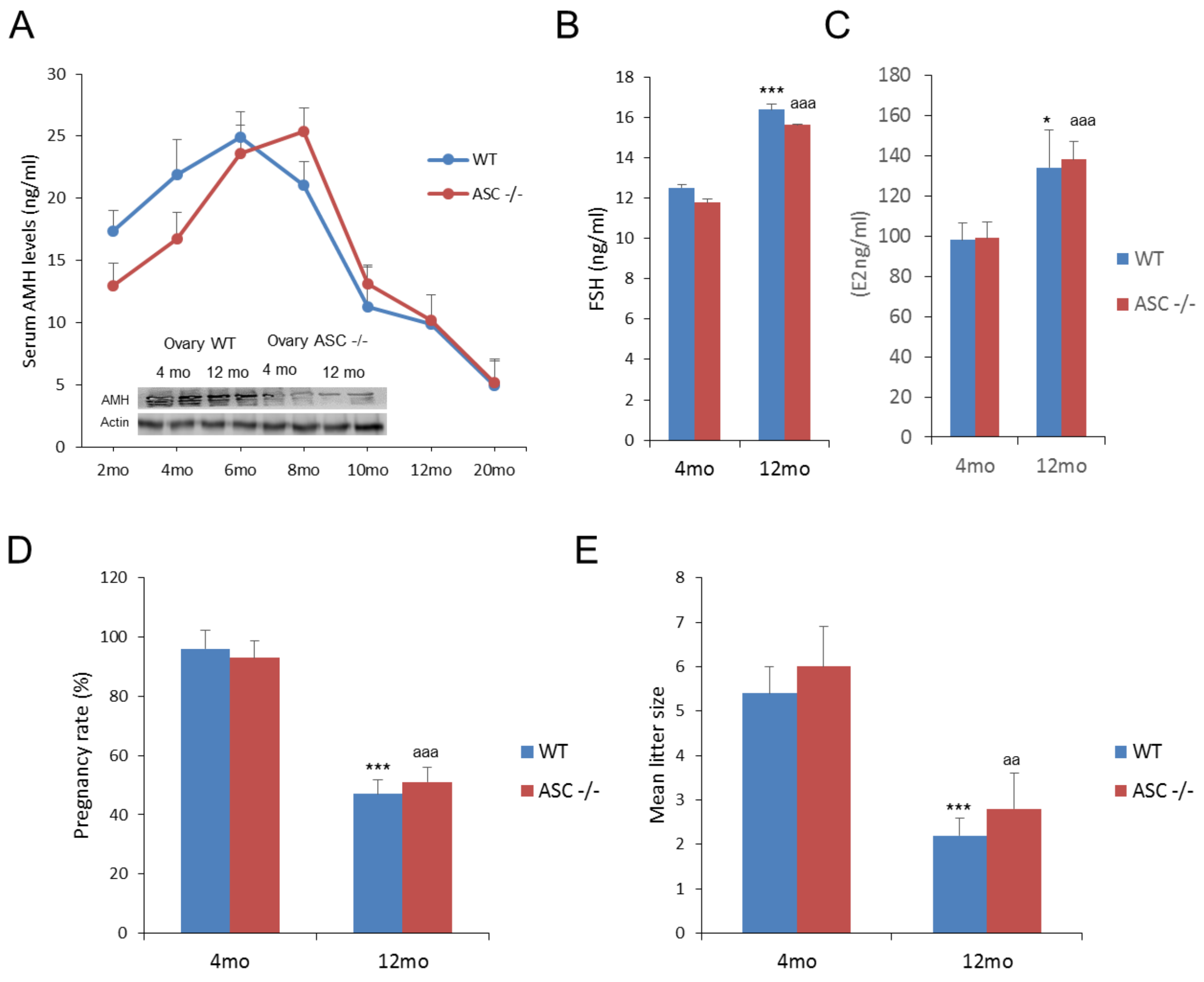


Supplementary Figure 3. (A) Mean serum AMH levels to evaluate the progression of ovarian reserve during aging in WT mice (blue) compared with ASC −/− mice (red) and ovarian AMH protein level. (B and C) Analysis of serum concen trations of FSH, and E2 as measured by ELISA. Pregnancy rate (D) and mean litter size (E) in aged WT and ASC -/- mice. All data are presented as means ± SD, n = 10 mice; *P < 0.05, **P < 0.005, ***P < 0.001 4mo *vs* 12mo WT mice; $^{a}$P < 0.05, $^{aa}$P < 0.005, $^{aaa}$P < 0.001 4mo *vs* 12mo ASC-/- mice

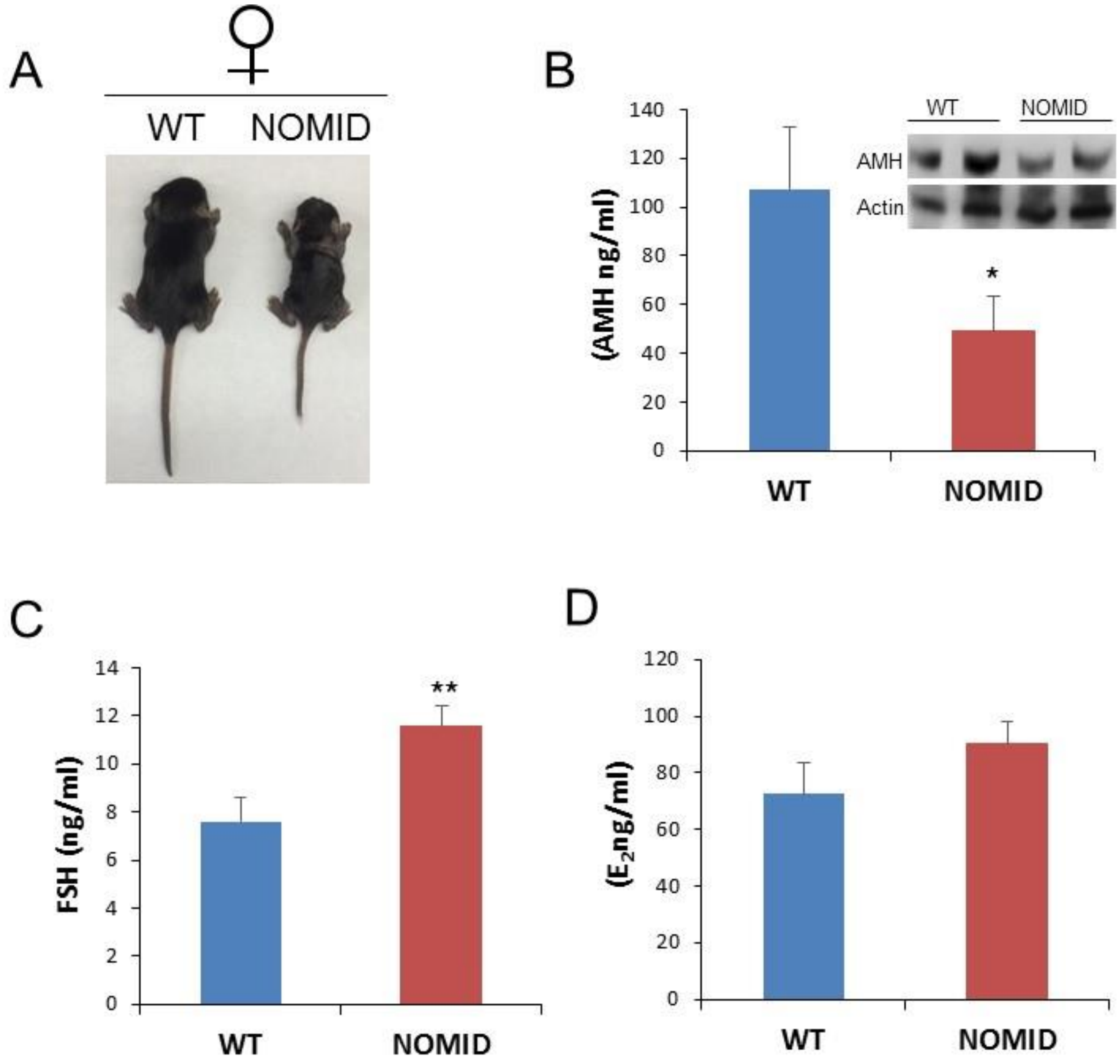


Supplementary Figure 4. (A) Representative photographs of 2-weeks-old NOMID and WT mice. (B) Mean serum AMH levels by ELISA and ovarian AMH protein levels (top) measured by WB, FSH (C) and E2 (D) serum levels in female WT compared with NOMID mice. Dara are presented as means ± SD. **P < 0.005, *P < 0.05 WT *vs* NOMID mice. n=5 mice per group.

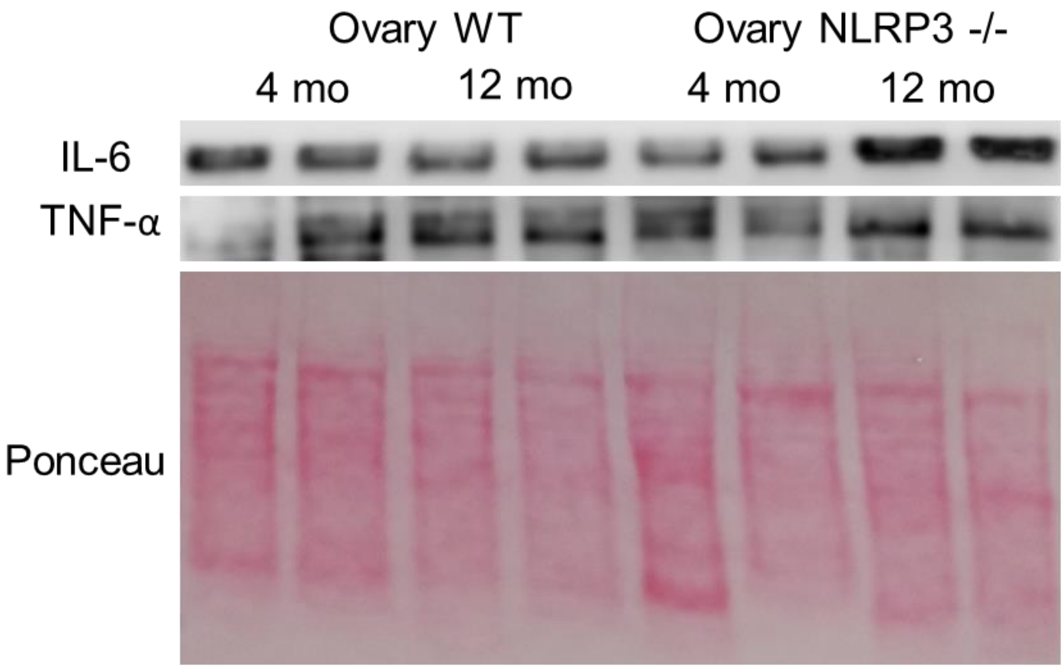


Supplementary Figure 5. Western blot analysis showing the protein expression of senescence markers IL-6 and TNF-α in the ovary of 4mo and 12mo NLRP3 -/- mice compared with WT. n= 4 mice per group and age

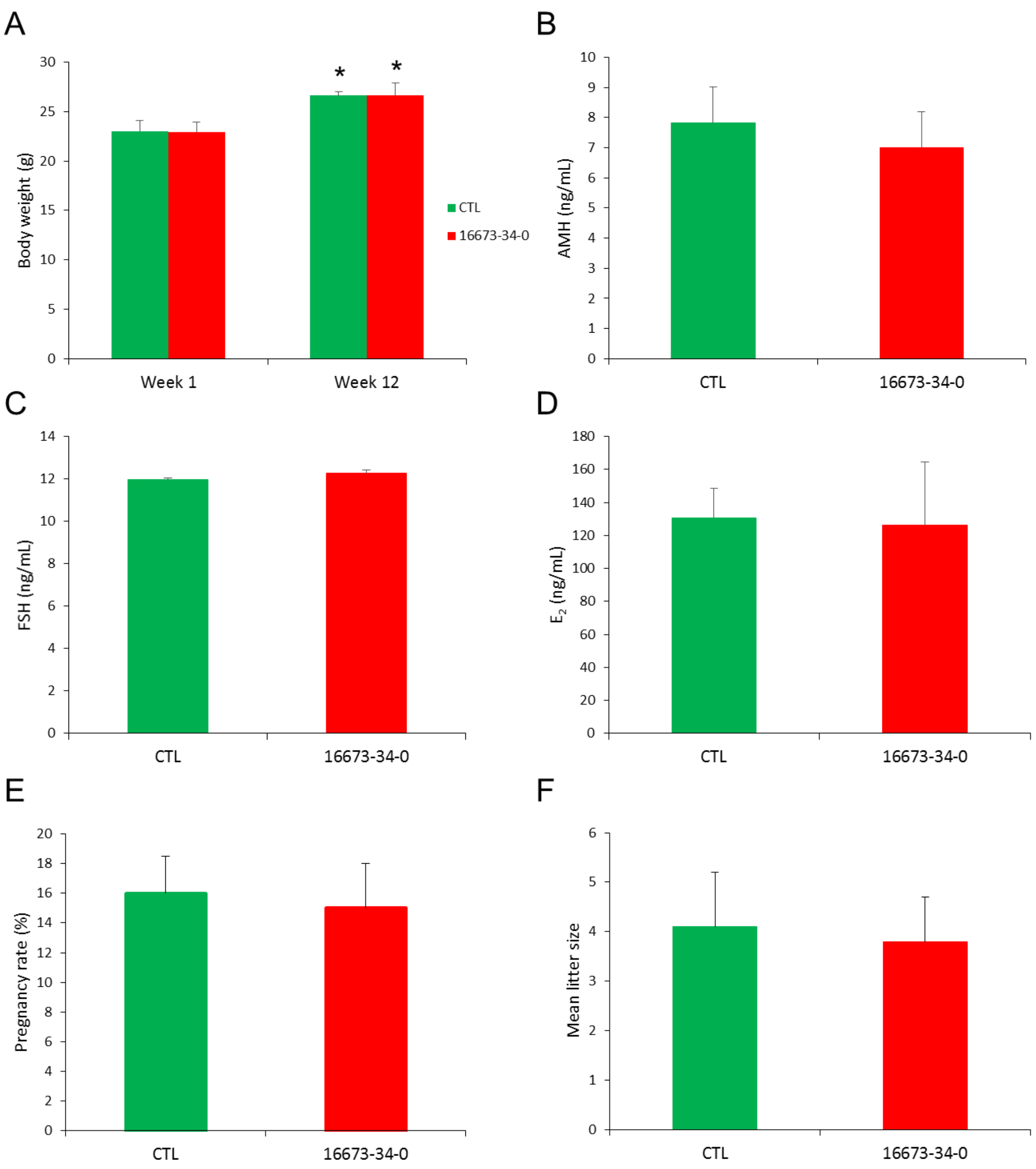


Supplementary Figure 6. CTL (vehicle) and 16673-34-0 treated female mice. (A) Body weights (B-D) Analysis of serum concen trations of AMH, FSH, and E2 as measured by ELISA. Pregnancy rate (E) and mean litter size (F) in vehicle and 16673-34-0 treated female mice. All data are presented as means ± SD, n= 6 mice; *P < 0.05, 1 week *vs* 12 weeks.
